# TAF-ChIP: An ultra-low input approach for genome wide chromatin immunoprecipitation assay

**DOI:** 10.1101/299727

**Authors:** Junaid Akhtar, Piyush More, Steffen Albrecht, Federico Marini, Waldemar Kaiser, Apurva Kulkarni, Leszek Wojnowski, Jean-Fred Fontaine, Miguel A. Andrade-Navarro, Marion Silies, Christian Berger

**Affiliations:** Institute for Developmental Biology and Neurobiology, University of Mainz, 55128 Mainz, Germany; Department of Pharmacology, University Medical Center, Johannes Gutenberg University of Mainz, 55131 Mainz, Germany; Center for Thrombosis and Hemostasis Mainz (CTH), 55131, Mainz, Germany; Institute of Medical Biostatistics, Epidemiology and Informatics (IMBEI), 55131 Mainz, Germany; Indian Institute of Science Education and Research (IISER) Pune, Maharashtra 411008, India; Faculty of Biology, Johannes Gutenberg University Mainz, Hans-Dieter-Hüsch-Weg 15, 55128, Mainz, Germany

## Abstract

Chromatin immunoprecipitation (ChIP) followed by next generation sequencing (ChIP-Seq) is a powerful technique to study transcriptional regulation. However, the requirement of millions of cells to generate results with high signal-to-noise ratio precludes it in the study of small cell populations. Here, we present a Tagmentation-Assisted Fragmentation ChIP (TAF-ChIP) and sequencing method to generate high-quality histone profiles from low cell numbers. The data obtained from the TAF-ChIP approach is amenable to standard tools for ChIP-Seq analysis, owing to its high signal-to-noise ratio. The epigenetic profiles from TAF-ChIP approach showed high agreement with conventional ChIP-Seq datasets, thereby underlining the utility of this approach.

## Introduction

Chromatin immunoprecipitation coupled with next generation sequencing (ChIP-Seq) is a powerful and unbiased approach to study genome-wide DNA-protein interactions and epigenetic modifications [1]. However, the prerequisite of huge starting material (millions of cells) limits its utility in studying rare cell types [2]. First, sonication; the by far most popular method for fragmentation in ChIP-Seq experiments, can destroy the epitope used for immunoprecipitation especially when the material is limited [3]. The alternative approach of micrococcal nuclease based digestion (MNase) is hard to control in its efficacy and saturation, and it also shows some degree of sequence dependent biases [4–6]. Second, the addition of sequencing adaptors for the generation of final libraries involves steps where the limitation of ligation and loss of material during purification steps can result in libraries with low-complexity.

Recently, there have been several attempts to adapt ChIP-Seq protocols to address these limitations in order to apply them to samples with low number of cells [7, 8]. One such method, called FARP-ChIP, used non-target cells for protection during sonication. To prevent the loss of DNA during library preparation, a biotinylated synthetic DNA (biotin-DNA) is used as a carrier DNA. The approach was successfully implemented to obtain the epigenetic profile from samples of 500 mESC cells. However, it required deep sequencing runs (approximately 100 million reads) and the number of reads mapping to the DNA of the target cell type was low (~16%), which makes this method less feasible for many applications and also cost intensive. Some other recent methods used prior ligation of barcoded adaptors to the chromatin digested by MNase, followed by a computational de-multiplexing strategy to obtain profiles from samples of low cell numbers [9]. The barcoding strategy was shown to dramatically reduce the number of cells required for each profile, and also can remove the biases arising from different chromatin preparations. However, the method still initially requires samples of 10,000-100,000 cells as starting material. Another approach, micro-scale μChIP-Seq, was used to generate the profile from samples of 500 cells. However, the method is a scaled down version of the conventional ChIP-Seq approach with samples subdivided at the level of immunoprecipitation [10]. The method ChIPmentation, uses Tn5 transposon mediated tagmentation for preparation of libraries as an alternative to the ligation based library preparation methods [11]. This reduces the hands-on time for library preparation and input requirements. However, this approach uses sonication for fragmenting the chromatin prior to immunoprecipitation. Moreover, this method still employs a large batch preparation of chromatin, and uses subsequent splitting of the sample to generate the profile from samples of 10,000 cells. Recently, the CUT&RUN approach was implemented to generate profiles from samples of 100 cells using antibody-targeted micrococcal nuclease [12]. The released and captured DNA was used to generate Illumina compatible libraries.

Here we describe an alternative approach for ChIP that uses Tagmentation-Assisted Fragmentation of chromatin (TAF-ChIP) with hyperactive Tn5 transposase from Illumina. The method employs limited sonication power only for nuclear lysis, and uses Tn5 activity for chromatin fragmentation. We have used this approach to generate high quality datasets from as few as 100 human and 1000 *Drosophila* cells. This approach has minimal hands-on time and does not involve labor-intensive library preparation workflow. Furthermore, it could be easily implemented to any type of cells. Comparisons of TAF-ChIP results to ENCODE datasets, CUT&RUN, and conventional ChIP-Seq performed in identical cell types demonstrates the utility of this approach. We expect our approach to be especially useful in conditions where the amount of sample is the limiting factor, such as material isolated from animals and clinical samples.

## Results

### Method Overview

There are two challenging steps in generating high quality ChIP-Seq datasets from samples with a very low number of cells. First, the fragmentation of chromatin without compromising the integrity of the associated proteins. Second, the generation of Illumina-compatible sequencing libraries, which requires the purified DNA to undergo multiple manipulation steps, namely end-repair, ligation of the sequencing adaptors, and PCR amplification. These steps also require beads-based purification of non-amplified DNA, where any potential loss of DNA can severely compromise the completion of successful libraries, especially when the starting amount of DNA is low. Furthermore, the intermediate steps can also be source of variability.

To overcome these limitations, we employed tagmentation as a tool to fragment the DNA. Tn5 mediated tagmentation had been previously used for the addition of sequencing adaptors on immunoprecipitated material, when preparing ChIP libraries and genomic DNA libraries.

Here, we instead used Tn5 activity to fragment the intact chromatin during immunoprecipitation. This approach has two major advantages. First, there is no need to fragment the chromatin before immunoprecipitation. Therefore, this strategy prevents potential loss of DNA-protein interactions during fragmentation, especially when compared to sonication. Furthermore, sonication is extremely variable between different machines, even if they are of the same specifications. Second, our tagmentation reactions employ the hyperactive Tn5 transposomes that are preloaded with sequencing adaptors [13]. Thus, after proteinase K inactivation, the immunoprecipitated material can be directly PCR amplified. This results in a one-step DNA library generation, which overcomes the limitation in efficiency of ligation and also avoids intermediate purification steps, thereby preventing loss of material [14]. After PCR amplification, the amplified libraries are bead-purified.

### Application of the TAF-ChIP approach on sorted *Drosophila* NSCs and human K562 cells

For TAF-ChIP samples, the cells were directly sorted into RIPA buffer owing to the low FACS sheath fluid volume, and directly preceded to nuclear lysis with low energy sonication. The low energy sonication employed here did not result in any visible fragmentation of chromatin. The non-fragmented chromatin was subjected to immunoprecipitation and tagmentation. After tagmentation, enzymes as well as background regions were washed away with subsequent high stringency washes. DNA was purified and PCR-amplified to generate Illumina compatible DNA libraries (see methods section for further details) (Figure 1 and Supplementary Figure 1A). For conventional ChIP-Seq samples, cells from *Drosophila* larval brain were sorted, pelleted and resuspended in RIPA buffer, as described earlier [15]. Upon immunoprecipitation with specific antibodies, the DNA was extracted and converted into DNA libraries (Supplementary Figure 1B). For the purpose of this study, we used two different type of starting material: type II neural stem cells (NSCs) from *Drosophila* larval brain and human K562 cells, a human immortalized myelogenous leukemia line. We used formaldehyde to fix freshly dissected *Drosophila* larval brains or harvested K562 cells. The dissected *Drosophila* larval brains expressed a GFP-tagged Deadpan (Dpn) protein under the control of its endogenous enhancer, which is a transcription factor only present in neural stem cells in the brain. This GFP was used to sort NSCs from wild type larvae, as described earlier [16].

**Figure 1.**
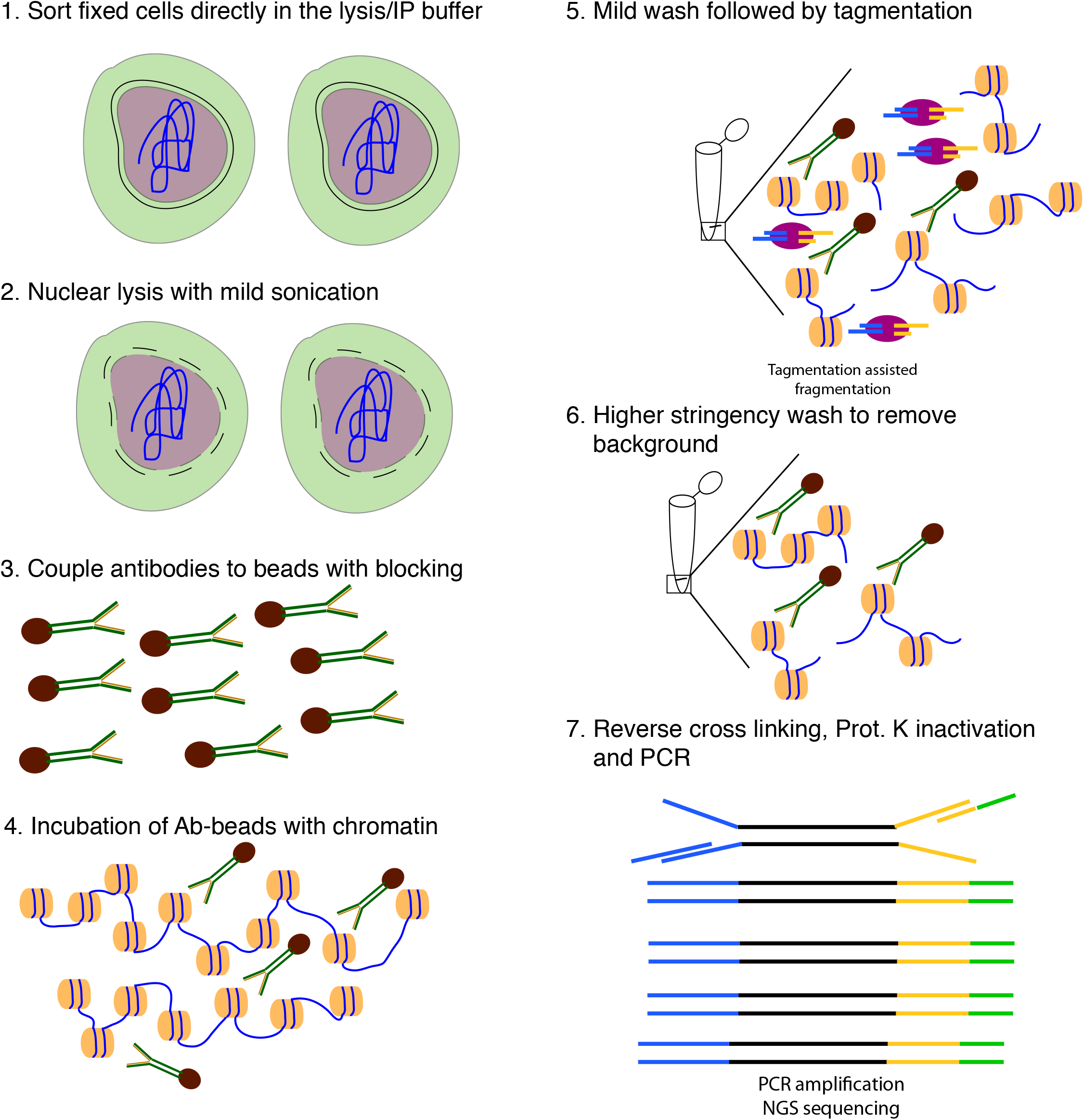
Schematic overview of TAF-ChIP approach. **(1)** Formaldehyde fixed cells were directly sorted into RIPA buffer (see methods for details). **(2)** Cells were briefly sonicated at low intensity to break open the nuclei. **(3)** Antibodies were coupled to magnetic beads in the presence of blocking reagents. **(4)** Antibody coupled beads were added to the cell lysate and incubated overnight at 4°C. **(5)** The tagmentation reaction was performed after initial washes with low salt IP buffer and homemade tagmentation buffer. **(6)** The tagmentation reaction and the background regions (not anchored by antibody interaction) were washed away with subsequent high stringency washes. **(7)** The proteinase K was heat-inactivated and the material was PCR-amplified without purification.

FACS-sorting of wild type NSCs is not applicable to obtain the ~1 million cells necessary to generate a conventional ChIP-Seq dataset, as one Drosophila brains consist of approximately 400 NSCs only. Thus, in order to compare the TAF-ChIP with the conventional ChIP-Seq protocol, we used the Gal4/UAS binary expression system to express a constitutively active Notch protein (*Notch^intra^*) in all type II NSCs (UAS/GAL4 system; *wor*-Gal4; *ase*-Gal80 fly line), also expressing UAS-CD8-GFP [17]. The expression of constitutively active *Notch^intra^* protein results in a massive over-proliferation of cells with the properties of type II NSCs amenable to cell sorting for conventional ChIP-Seq [18].

We sorted type II NSCs from this line with identical settings as above, for TAF-ChIP (1000 cells) as well as for conventional ChIP-Seq (1.2 million cells). For obtaining 100 K562 cells, we stained the cells with Hoechst dye and used FACS for collecting samples with the precise number of cells. To benchmark our TAF-ChIP data sets from K562 cells, we used publicly available datasets from the ENCODE project [19, 20].

The Tn5 tagmentation is preferably done in the open chromatin region due to higher accessibility (which is the basis of the ATAC-Seq approach), and thus these regions can get over-represented [21]. To distinguish from this scenario and to get a better estimate of background signal, we also performed TAF-ChIP experiments with histone H3.

### Detailed evaluation of TAF-ChIP

To investigate in detail the performance of TAF-ChIP against both the conventional ChIP-seq and the recently described CUT&RUN low amount method, we used receiver-operating characteristic (ROC) curves and precision-recall (PR) curves [7]. Towards this goal, we compared the peaks in K562 cells for the TAF-ChIP datasets, conventional ENCODE datasets and CUT&RUN datasets for 100, 3000, and 6000 cells at various FDR cutoffs and using the *replicated peaks* of the conventional ENCODE dataset as reference [12, 19]. K652 curves were calculated by mapping peaks to 5kb non-overlapping genomics windows. Similarly, we also compared peaks for TAF-ChIP and for conventional ChIP-seq datasets from *Drosophila* UAS-derived NSCs at various FDR cutoffs and using the first replicate of the conventional ChIP-seq dataset as reference. NSC curves were calculated by mapping peaks to 1kb non-overlapping genomic windows. The peaks were always obtained with MACS2 peak calling algorithm using either input (conventional ChIP-Seq) or H3 datasets (TAF-ChIP) as controls.

For human K562 cells, both the ROC curves and the precision recall curves showed that the 100 cells TAF-ChIP dataset was comparable to the reference ENCODE replicate, as well as to 3000 and 6000 cells CUT&RUN datasets, outperforming the 100-cells CUT&RUN dataset (Figure 2A and 2B). Only ~500 peaks were called at 5% FDR for the 100-cells CUT&RUN dataset. This could be due to high occurrence of noise in the 100-cells CUT&RUN dataset, which can be observed in the genome browser profile (Supplementary Figure 1C). The CUT&RUN method on 100 cells was not able to recall more than 75% of the reference even though the peak calling parameters had no restrictions.

**Figure 2.**
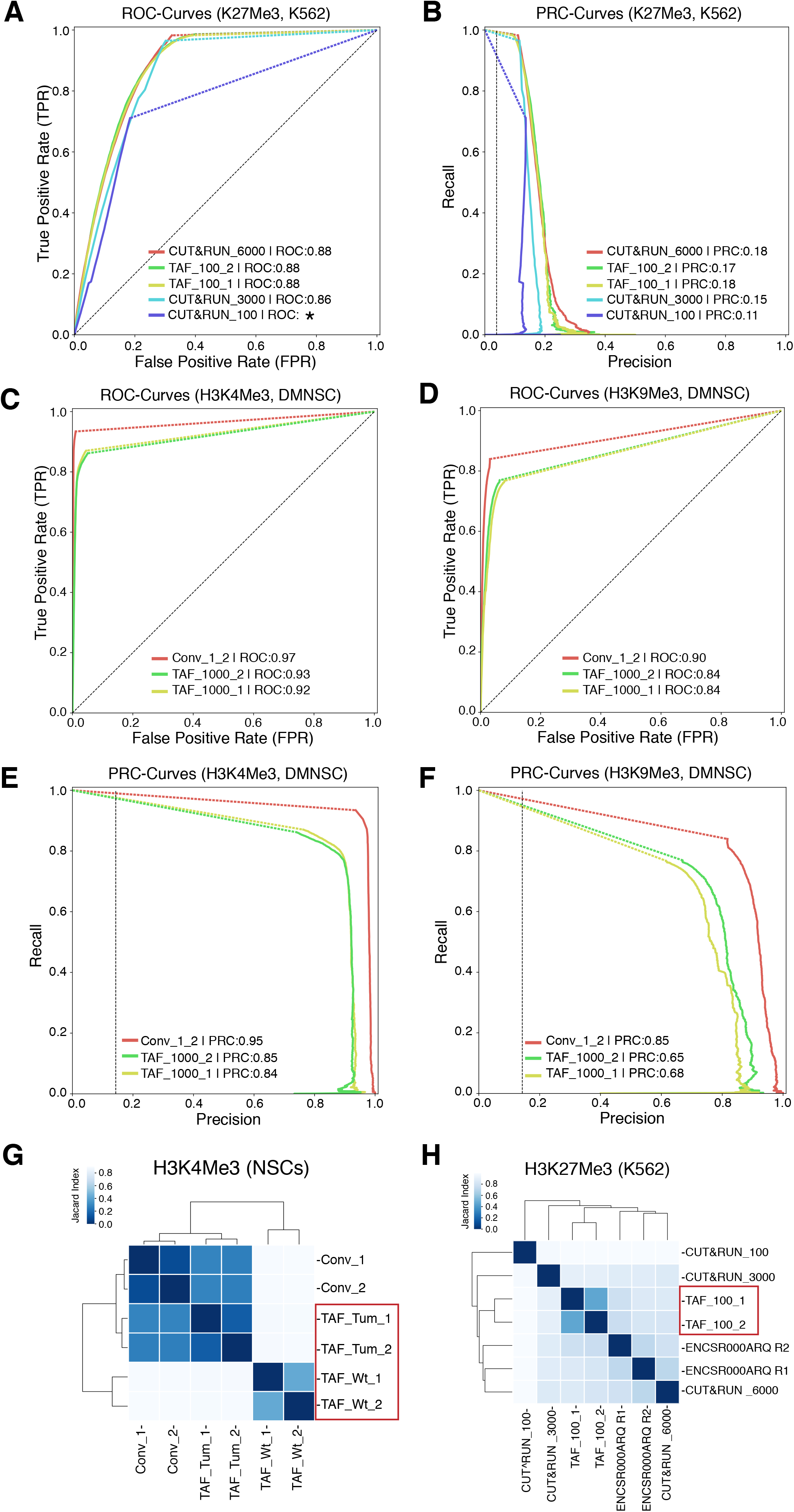
Comparison of TAF-ChIP with conventional ChIP-Seq and with the CUT&RUN low amount method. **(A)** ROC curves of TAF-ChIP and CUT&RUN for H3K27Me3 in K562 cells. The ROC curves were plotted using as reference replicated peaks of the conventional ChIP-Seq ENCODE dataset selected at 5% FDR cutoff (downloaded from the ENCODE database). No FDR cutoff was used to define peaks for TAF-ChIP replicates and the CUT&RUN datasets with MACS2. Peaks were mapped to 5kb non-overlapping genomic windows to calculate true positive rate or recall, false positive rate and precision for a changing p-Value threshold. Area under the curve (AUC) is indicated in the legend in decreasing order, and the * indicates the failure to faithfully calculate the AUC. **(B)** Precision recall curve for TAF-ChIP and CUT&RUN datasets for H3K27Me3 in K562 cells. **(C, D)** Receiver operating characteristic (ROC) curves of TAF-ChIP and conventional ChIP-Seq in *Drosophila* NSCs. The ROC curves for H3K4Me3 **(C)** and H3K9me3 **(D)** were plotted using as reference peaks of the first conventional ChIP-Seq replicate selected at 5% FDR cutoff. No FDR cutoff was used to define peaks for TAF-ChIP replicates and the second conventional ChIP-Seq replicate. Peaks were mapped to 1kb non-overlapping genomic windows to calculate true positive rate or recall, false positive rate, and precision. Area under the curve (AUC) is indicated in the legend in decreasing order. **(E, F)** Precision recall curve for TAF-ChIP and conventional ChIP-Seq in Drosophila NSCs. Using same references and data as above, precision-recall curves were plotted for H3K4Me3 (E) and H3k9Me3 (F). **(G)** Comparison of the genomic window sets for *Drosophila* brain-derived wt NSCs analysed for H3K4Me3 binding by TAF-ChIP (TAF_Wt), and *Drosophila* tumour-derived NSCs analysed by TAF-ChIP or conventional ChIP-Seq (TAF_Tum and Conv). The TAF-ChIP samples are highlighted by a red rectangular box. The heatmap indicates pairwise similarity according to the Jaccard Index. Axes show results of hierarchical clustering. **(H)** Similar comparison and representation as in (G) for K562 cells H3K27Me3 binding assayed with TAF using a 100-cell sample (TAF_100, highlighted with a red rectangular box), with CUT&RUN method for 100, 3000 and 6000-cell samples and with conventional ChIP-Seq (ENCODE) [12, 19].

For *Drosophila* NSCs, the ROC and PRC curves showed that our TAF-ChIP approach has a comparable performance to the inter-replicate results of conventional ChIP-Seq (Figure 2C-F).

We next compared the datasets by hierarchical clustering using a similarity measure based on the Jaccard Index calculated on sets of genomic windows from peaks defined at 5% FDR. The conventional H3K4Me3 ChIP-Seq datasets from *Drosophila* NSCs (tumor-derived) clustered together with H3K4Me3 TAF-ChIP datasets from tumor NSCs rather than with wild type NSCs (Figure 2G). For K562 cells, the H3K27Me3 TAF–ChIP datasets clustered together with the corresponding ENCODE dataset and with CUT&RUN datasets from higher cell numbers (Figure 2H). Consistent with our ROC curve and PRC curve analysis, the 100-cell CUT&RUN dataset showed lower similarity to the rest of the datasets.

We also plotted the hierarchical clustering for H3K9Me3 and H3K27Me3 with other histone ChIP-Seq datasets included as control. The TAF-ChIP datasets always clustered together with their corresponding ENCODE datasets rather than with unrelated histone ChIP-Seq (Supplementary Figures 1D and 1E). The TAF-ChIP dataset for H3K9Me3 from *Drosophila* NSCs (tumor) also clustered together with conventional ChIP-Seq performed in identical NSCs (Supplementary Figure 1F).

### Comparison of 100 cells TAF-ChIP to ENCODE dataset

To further test the applicability of TAF-ChIP we next used corresponding conventional ChIP-Seq datasets from the ENCODE project for benchmarking.

The H3K27Me3 TAF-ChIP and H3K9Me3 TAF-ChIP from samples of 100 cells showed similar profiles when compared with the corresponding ENCODE datasets, as visualized through genome browser tracks (Figure 3A and 3B), and also had good agreement between the replicates when Pearson’s correlation coefficient was calculated using average signal in each 2kb non-overlapping genomic window (Supplementary Figures 2A and 2B). The metagene profile for H3K27Me3 and H3K9Me3 showed decrease at the TSSs and higher signal on the gene body, similar to the profile obtained with the ENCODE dataset (Figures 3C and 3D). We used the MACS2 peaks calling algorithm for identifying the peaks in both TAF-ChIP and ENCODE datasets, with identical parameters. The corresponding input samples, fragmented input control for ENCODE and H3 TAF for TAF-ChIP, were used as control for peak calling. The annotation of peaks identified in the TAF-ChIP dataset and in the corresponding one from ENCODE showed similarity in distribution of overlapping genomic features, for both H3K27Me3 and H3K9Me3 datasets (Figure 3E). The overlap between the peaks called for ENCODE and TAF-ChIP was 50% for H3K27Me3 and 56% for H3K9Me3 (Figure 3F). Next, we divided the peaks into 10 different quantiles according to FDR, with quantile 1 associated with the lowest FDR peaks and quantile 10 associated with the highest. The FDR quantile recovery analysis for H3K27Me3 and H3K9Me3 peaks compared to replicated peaks of ENCODE and was higher for lower FDR quantiles, at around 60% and 70%, respectively (Supplementary Figure 2C and 2D). The fraction of reads in peaks called with TAF-ChIP had also similar distribution profile when compared to the ENCODE ChIP-Seq. However, the level of this enrichment was smaller for TAF-ChIP (Figure 3G). Nonetheless, the heatmaps generated for all the peaks identified in ENCODE ChIP-Seq datasets and sorted according to the intensity in the ENCODE ChIP-Seq, showed profiles similar and comparable to TAF-ChIP datasets from 100 K562 cells (Figure 3H).

**Figure 3.**
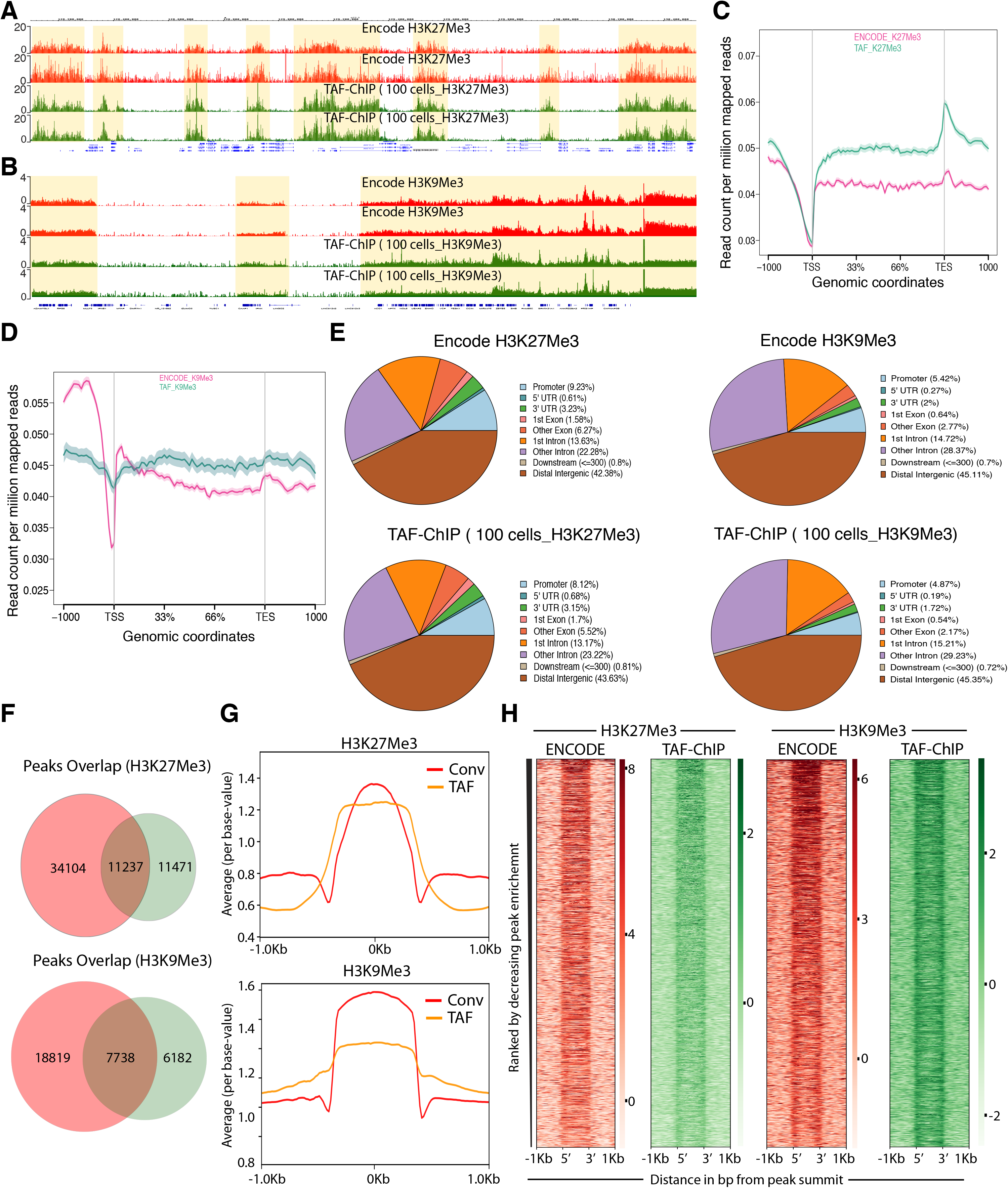
TAF-ChIP results from 100 K562 cells are comparable to conventional Encode ChIP-Seq datasets. **(A, B)** Genome browser track example of H3K27Me3 and H3K9Me3 (A and B, respectively) ChIP performed in 100 FACS sorted K562 cells with TAF-ChIP approach and corresponding K562 conventional ChIP-Seq datasets from the ENCODE project in duplicates, as indicated in the labels. The label below the tracks shows the gene model, and the y-axis represents normalized read density in reads per million (rpm). The enriched regions are highlighted with shaded box. **(C, D)** Metagene profiles of H3K27Me3 and H3K9Me3 (C and D, respectively) with standard error to the mean for all the genes, −1000bp upstream of transcription start sites (TSS) and +1000bp downstream of transcription end sites (TES). Read counts per million of mapped reads is shown on the y-axis, while the x-axis depicts genomic coordinates. **(E)** Genomic distribution of annotated peaks obtained from the ENCODE datasets and TAF-ChIP (100 K562 cells), for indicated histone marks. Note the majority of H3K27Me3 and H3K9Me3 peaks are at the intergenic regions, consistent with the expectation. **(F)** Overlap between the peaks identified from the ENCODE and TAF-ChIP datasets, for the indicated histone modifications (see methods section for further details). **(G)** Average profile of TAF-ChIP and corresponding ENCODE ChIP-Seq centered at the peaks for the indicated histone modifications. The y-axis depicts average per base-value into the peaks while x-axis depicts genomic coordinates centered at the peaks. **(H)** Distributions of reads at gene locations of indicated histone modifications from ENCODE ChIP-Seq and TAF-ChIP method, centered at the peaks (−1kb to +1kb). Rows indicate all the peaks and are sorted by decreasing affinities in the ENCODE ChIP-Seq data sets. The color labels to the right indicate the level of enrichment.

### TAF-ChIP performed on *Drosophila* NSCs shows high agreement with conventional ChIP-Seq

To compare TAF-ChIP to conventional ChIP-Seq, we analysed both H3K4Me3 and H3K9Me3 histone marks, from identical cell types, as described above. The TAF-ChIP generated datasets showed similar signal-to-noise ratio when compared with corresponding conventional ChIP-Seq datasets, as visualized through genome browser tracks (Figures 4A and 4B). The TAF-ChIP data also showed high degree of mappability and low level of sequence duplication. The uniquely mapped reads for H3K4Me3 samples were at ~80%. The unique mapping rate for H3K9Me3 was lower at ~60%, yet this can be expected due to prevalence of this mark at repeat elements and transposons (Table S3A). The replicates also showed good concordance between themselves when Pearson’s correlation coefficient was calculated using average signal in each 500bp non-overlapping genomic windows (Supplementary Figures 3B and 3C). The metagene profile for H3K4Me3 normalized to H3 and IgG control showed higher signal at the TSSs, consistent with the higher enrichment of this mark at the promoters (Figure 4C). On the other hand, the metagene profile for H3K9Me3 showed higher enrichment over gene body (data not shown). Furthermore, the qPCR analysis of TAF-ChIP and conventional ChIP for H3K4Me3 showed comparable enrichment for enriched loci, as well as similar level of background at non-enriched locus (Supplementary Figure 3D). Next, we used the MACS2 software [21] to identify peaks in the TAF-ChIP and in conventional ChIP-Seq datasets. The fragmented input control and H3 TAF-ChIP datasets were used as input control for conventional ChIP-Seq and TAF-ChIP datasets for peak calling, respectively. The deposition of H3K9Me3 was mostly on intergenic regions, therefore we utilized peak coordinates to generate the normalized metagene profile (Figure 4D). The annotation of peaks obtained from TAF-ChIP and ChIP-Seq showed a higher degree of similarity for the H3K4Me3 mark than for the H3K9Me3 mark, the latter displaying more overlap to promoters and less to intergenic regions in conventional ChIP-Seq (Figure 4E). Nevertheless, consistent with the expectation, the large fraction of H3K4Me3 peaks was at the promoters whereas the majority of H3K9Me3 peaks were at the distal intergenic regions. Next, we calculated the overlap between the peaks called for conventional ChIP-Seq and TAF-ChIP datasets using the ChIPpeakAnno package from bioconductor [22]. The peaks called for H3K4Me3 showed 85% overlap between the conventional and TAF-ChIP approaches at 5% FDR. The peaks called at 5% FDR for H3K9Me3 had 68% of overlap between the conventional and TAF-ChIP approach (Figure 4F).

**Figure 4.**
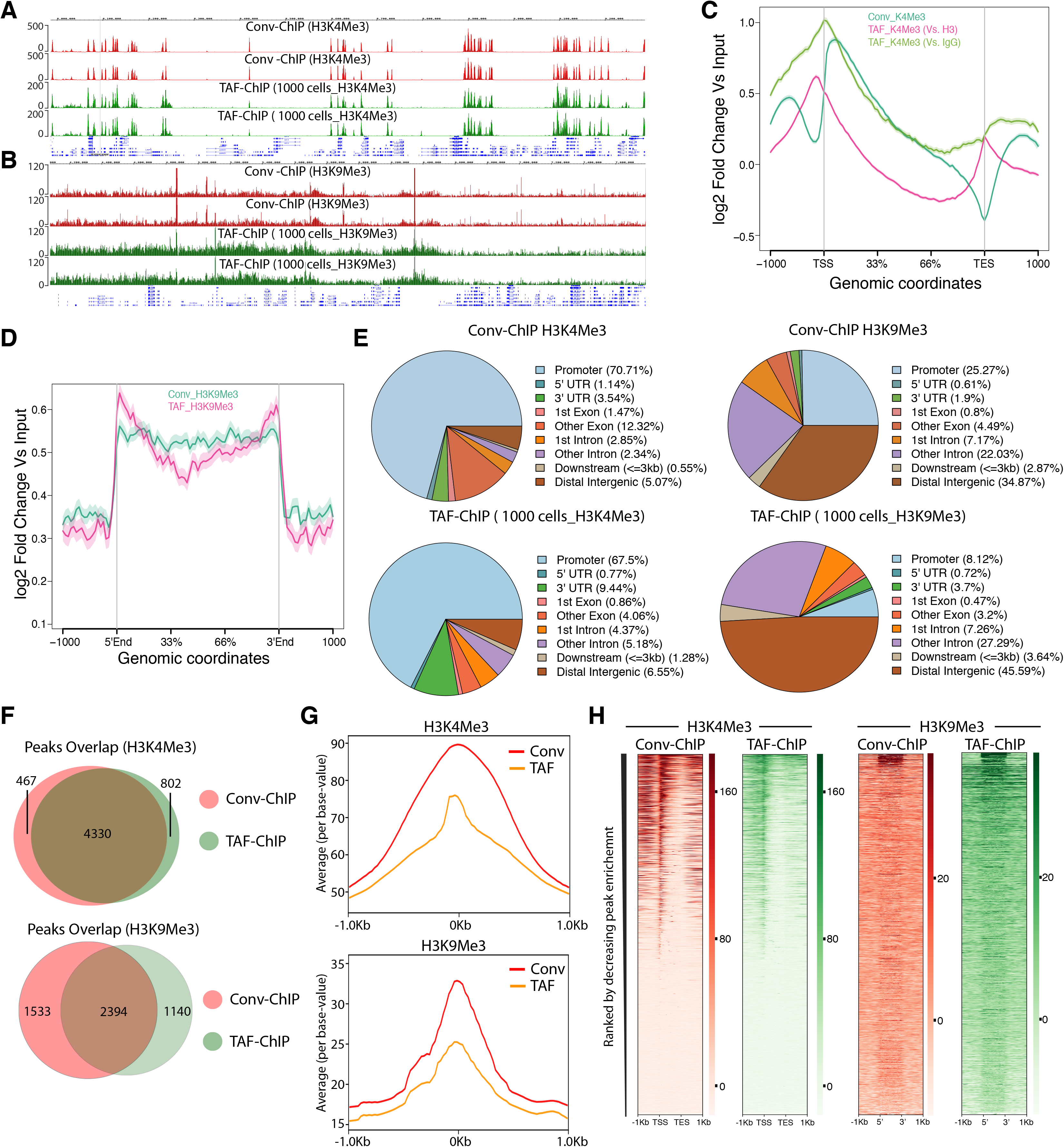
TAF-ChIP results from low number of NSCs are comparable to conventional ChIP-Seq (Conv-ChIP). **(A, B)** Genome browser track example of H3K4Me3 and H3K9Me3 ChIP (panel A and B, respectively) performed in FACS sorted NSCs with conventional ChIP-Seq (1.2 million cells) and TAF-ChIP (1000 cells), as indicated by the labels. The label below the tracks shows the gene model, and the y-axis represents normalized read density in reads per million (rpm). **(C)** Metagene profiles of H3K4Me3 with standard error to the mean for all genes, - 1000bp upstream of transcription start sites (TSSs) and +1000bp downstream of transcription end sites (TESs). Log2 fold changes against input controls are shown on the y-axis, while the x-axis depicts genomic coordinates. **(D)** Metagene profiles of H3K9Me3 with standard error to the mean for enriched regions, −1000bp upstream and +1000bp downstream of peaks. Log2 fold changes against input control are shown on the y-axis, while the x-axis depicts genomic coordinates. **(E)** Distribution of annotated peaks obtained from conventional ChIP-Seq and TAF-ChIP, for indicated histone marks. Note that the majority of H3K4Me3 and H3K9Me3 peaks are at the promoters and at the intergenic regions, respectively, consistent with the expectation. **(F)** Overlap between the peaks identified from conventional ChIP-Seq and TAF-ChIP datasets, for the indicated histone modifications. MACS2 software with identical parameters (see methods for details) was used to identify the peaks against the respective input controls, and those present in both replicates were considered for the comparison. **(G)** Average profile of TAF-ChIP and conventional ChIP-Seq centered at the peaks for the indicated histone modifications. The y-axis depicts average per base-value into the peaks while the x-axis depicts genomic coordinates centered at the peaks. **(H)** Distributions of reads at gene locations of indicated histone modifications from conventional ChIP-Seq and TAF-ChIP. Rows indicate all the peaks and are sorted by decreasing affinities in the conventional ChIP-Seq data sets. The color labels to the right indicate the level of enrichment.

Next, we performed the peak recovery in different FDR quantiles, as explained before for K562 datasets. Using one H3K4me3 conventional ChIP-seq replicate as reference, TAF-ChIP recalled ~99% of the peaks until quantile 6, and was comparable to the other replicate of the conventional ChIP-Seq (Supplementary Figure 3E). The relationship between recall and FDR was very weak for H3K9Me3, however it was still similar to conventional ChIP-Seq (Supplementary Figure 3F). The read distribution at the peaks still showed enrichment for TAF-ChIP, albeit to a lower level when compared to conventional ChIP-Seq datasets (Figure 4G). The analysis for saturation of peak recall showed higher recall of peaks for H3K4Me3 at shallow sequencing depth, whereas for H3K9Me3 the number of recalled peaks continued to increase with increasing sequencing depths (Supplementary Figure 3G and 3H). This was consistent with the observed tendencies for point-source histone modifications (such as H3K4Me3) and histone modifications with broad domains of enrichments (such as H3K9Me3) [20]. The distributions of reads at genomic locations generated for TAF-ChIP and conventional ChIP-Seq datasets, and sorted according to the intensity in the conventional ChIP-Seq resulted in similar and comparable profiles (Figure 4H).

### TAF-ChIP gave consistent results with variable numbers of cells used as starting material

After establishing TAF-ChIP on low number of cells and its subsequent benchmarking against conventional ChIP-Seq performed in identical cells, we next assayed whether TAF-ChIP can give comparable results with similar resolution, when variable numbers of cells are used as starting material. Towards this goal, we resorted to use wild type NSCs from *Drosophila* brains. We sorted two samples containing 1000 and 5000 NSCs, respectively. The TAF-ChIP generated datasets from 1000 and 5000 NSCs resulted in nearly identical profiles, as visualized through genome browser tracks (Supplementary Figure 4A). The distributions of reads at genomic locations generated from 1000 NSCs and 5000 NSCs also showed comparable profiles (Supplementary Figure 4B). The read distribution in the peaks for samples with 1000 NSCs and 5000 NSCs were also comparable to each other (Supplementary Figure 4C). Altogether, these results suggest that TAF-ChIP is amenable to conditions when starting material is variable to few folds, and would produce similar results.

## Discussion

Here, we present an easy, tagmentation-assisted fragmentation ChIP (TAF-ChIP) and sequencing method to generate high-quality datasets from samples with low cell numbers. The workflow of TAF-ChIP contains fewer steps than conventional ChIP-Seq with minimum hands-on-time during library preparation, preventing loss of material and potential user introduced variability. Due to tagmentation during immunoprecipitation the cells can be directly sorted into the IP/lysis buffer. This eliminates the centrifugation step to collect the cells, which can also lead to potential loss of material. Also, unlike ATAC-Seq where intact nuclei are tagmented and partial tagmentation is used to study chromatin accessibility, our approach tagments after nuclear lysis [21]. The metagene profile of H3 TAF-ChIP dataset did not show any enrichment for TSS, suggesting our method resulted in tagmentation without any visible biases for open chromatin regions (Supplementary Figure 4D and 4E). Furthermore, we also showed the application of TAF-ChIP for both open chromatin marks such as H3K4Me3 as well as for repressive marks such as H3K9Me3 and H3K27Me3. TAF-ChIP is easier to implement than MNase based approaches, does not lead to over-digestion of chromatin, and results in one-step generation of Illumina compatible libraries. The TAF-ChIP approach is suitable for assaying factors where the chromatin association might be dependent on RNA, as Tn5 does not perturb the RNA intermediate. Also, the tagmentation does not show any sequence dependent biases, in contrast to other restriction-based protocols [6, 13]. Furthermore, the approach does not need any specialized equipment and thus can be implemented in a standard molecular biology lab.

We have used here the Tn5 transposase from Nextera XT DNA library kit; however, TAF-ChIP could be easily implemented with Tn5 loaded with different unique molecular indices [13]. This could be easily implemented in massively parallel TAF-ChIP Seq applications, and may even further decrease the required starting material. Moreover, as this approach can be used for various cell types, it could be also combined with a non-target cell type used as “spike-in” and DNA carrier.

We showed that TAF-ChIP datasets have signal-to-noise ratios that are comparable to conventional ChIP-seq datasets and thus are amenable to standard bioinformatics pipelines for ChIP-Seq analysis. Our evaluation of TAF-ChIP datasets showed results comparable to conventional ChIP-Seq and better than CUT&RUN, a comparable low amount method. For histone marks, we demonstrated the use of H3 TAF-ChIP as an input control for better background estimation. However, in some conditions and perturbation experiments, the distribution of H3 might have weak to strong biases. An alternative control for TAF-ChIP could be also immunoprecipitation with IgG from similar species (Figure 4C). Although the genome browser profiles obtained from a sample of 100 K562 cells showed slightly inferior signal to noise ratio compared to the conventional datasets from the ENCODE project (Figure 2A) yet the peaks identified were mostly overlapping, especially at lower FDRs. The peaks that were unique to either TAF-ChIP or to the conventional method still showed higher read coverage compared to randomly selected regions of comparable size (Supplementary Figure 5A-5D). This suggests that thresholding (based on FDR) implemented by the peak caller software might have hindered their identification in one of the datasets. Furthermore, we suspect that the signal to noise ratio can be improved by pooling the samples tagmented with different indices, prior to washes and following the demultiplexing strategy to obtain the data.

We have also been able to generate the profile of a RNA modifying enzyme, recently shown in vertebrates to associate with chromatin, by performing TAF-ChIP using 1000 cells from transgenically tagged *Drosophila* and antibody directed against the tag (unpublished results). We conclude that the only limiting factors defining the lowest cell number sample providing biologically meaningful TAF-ChIP results are the availability of a good antibody and a reasonable number of binding sites in the genome. We have shown that TAF-ChIP provides reliable datasets from samples of as low as 100 isolated cells without requiring prior isolation of nuclei and with an extremely easy and straightforward workflow; therefore, we expect that TAF-ChIP will be very useful when access to higher numbers of cells are limited.

## Author Contributions

J.A. and C.B. designed the experiments. J.A. performed the experiments. J.A, P.M, S.A, J.F and F.M. performed the bioinformatics analysis of the data. A.K. and W.K. provided the *Drosophila* NSC samples. J.A. wrote the paper with the help and feedback of rest of the authors.

## Acknowledgments

We thank the Bloomington *Drosophila* Stock Center for the fly lines used in this study. The EMBL Genomics Core Facility (Heidelberg, Germany), especially Vladimir Benes for all the sequencing runs. We thank Jean-Yves Roignant, Guillaume Junion, Joachim Urban, and Prasad Chitke for critically reading the manuscript. Research in the laboratory of C.B. is supported by the Deutsche Forschungsgemeinschaft (DFG) DFG BE 4728 1-1 and 3-1, and the University of Mainz. P.M., S.A. and A.K. thank the International PhD Programme (IPP) of the Institute of Molecular Biology, Mainz for supporting the PhD. The work of F.M. is supported by the German Federal Ministry of Education and Research (BMBF 01EO1003).

## Accession number

All the ChIP-Seq data generated in this study is submitted to the GEO database (GSE112633).

## Methods

### Antibodies

The following antibodies were used in this study. For H3K4Me3 ChIP, antibody from Abcam (Cat No-ab8580) was used. H3K9Me3 ChIP was performed using an antibody from Active Motif (Cat No-39161), H3K27Me3 (Active Motif, 39155) and H3 (Abcam, ab1791).

### Fixation and cell sorting from *Drosophila* larval brain

Briefly, required number of larval brains after 48h of larval hatching were dissected in PBS. After dissection, larval brains were fixed with 1% formaldehyde in PBS for 10 min at room temperature, followed by quenching of the fix with 125 mM glycine. The larval brains were dissociated and resuspended according to the previously established method [16]. The cells were then sorted on BD FACSAria™ according to the strength of GFP and size of the NSCs, resulting in a pure population of type II neural stem cells.

### Fixation and cell sorting of K562 cells

K562 cells cultured in RPMI medium (supplemented with 10% Fetal Bovine Serum), at 37°C and 5% CO_2_ were fixed for 10 min at room temperature with 1% formaldehyde. The crosslink was quenched with 125 mM glycine, and sorted on BD FACSAria™ cell sorter using Hoechst stain. A total of 100 K562 cells were directly sorted in RIPA 140 mM (10 mM Tris-Cl pH 8.0, 140 mM NaCl, 0.5 mM EDTA pH 8.0, 1% Triton-X 100 and 0.1% SDS).

### Conventional ChIP-Seq and library preparation

Fixed cells (1.2 million FACS sorted NSCs per replicate) were resuspended in 140 mM RIPA (10 mM Tris-Cl pH 8.0, 140 mM NaCl, 0.1 mM EDTA pH 8.0, 1% Triton X-100, and 0.1% SDS) and subjected to 14 cycles of sonication on a bioruptor (Diagnode), with 30 Secs “ON”/ “OFF” at high settings. After sonication, samples were centrifuged at 14,000 g for 10 min at 4°C and supernatant was transferred to a fresh tube. The extracts were incubated overnight with 2 μg of specific antibody at 4°C with head-over tail rotations. After overnight incubations, 20 μl of blocked Protein A and G Dynabeads were added to the tubes and further incubated for 3 hrs to capture the antibodies. The beads were separated with a magnetic rack and were washed as following; once with 140 mM RIPA (10 mM Tris-Cl pH 8.0, 140 mM NaCl, 0.1 mM EDTA pH 8.0, 1% Triton X-100, and 0.1% SDS), four times with 250 mM RIPA (10 mM Tris-Cl pH 8.0, 250 mM NaCl, 0.1mM EDTA pH 8.0, 1% Triton X-100, and 0.1% SDS) and twice with TE buffer pH 8.0 (10 mM Tris-Cl pH 8.0 and 0.1 mM EDTA pH 8.0). After the immunoprecipitation, samples were RNase-treated (NEB) and subjected to Proteinase K treatment for reversal of cross-links, 12 hrs at 37°C and at least 6 hrs at 65°C. The samples after proteinase K treatment were subjected to phenol chloroform extraction. After precipitating and pelleting, DNA was dissolved in 30 μl of TE buffer pH 8.0. The recovered DNA was converted into libraries using NebNext Ultra II DNA library preparation kit, following manufacturer’s protocol.

### TAF-ChIP and library preparation

Fixed cells were directly sorted in 240 μl of 140 mM RIPA (10 mM Tris-Cl pH 8.0, 140 mM NaCl, 0.1 mM EDTA pH 8.0, 1% Triton X-100, and 0.1% SDS), and sonicated with 3 cycles at low power settings for breaking the nuclei. 15 μl of Protein A and G Dynabeads were coupled to 1 μg of specific antibody in the blocking buffer (RIPA 140 mM supplemented with 0.2 mg/ml BSA, 0.05 mg/ml of glycogen and 0.2 mg/ml of yeast tRNA) for 3 hrs at 4°C. The predominantly unfragmented chromatin were centrifuged at 2000g for 10 min at 4°C, and the supernatant was transferred to the tube with blocked and antibody coupled beads. The samples can be also added to the tube with blocked and antibody coupled beads, without subjecting them to centrifugation. The samples were incubated at 4°C overnight with head over tail rotations. The samples were then washed twice briefly with 300 μl of homemade tagmentation buffer (20 mM Tris(hydroxymethyl)aminomethane pH 7.6; 10 mM MgCl_2_; 20% (vol/vol) dimethylformamide) using magnetic rack for beads separation. The washed beads were resupended in 20 μl of 1X tagmentation DNA buffer (Nextera XT Kit) containing 1 μl of Nextera DNA tagmentation enzyme and incubated at 37 °C for 40 min with constant shaking in a thermoblock at 500 rpm. Following the tagmentation, the beads were washed as following; once with 140 mM RIPA (10 mM Tris-Cl pH 8.0, 140 mM NaCl, 0.1 mM EDTA pH 8.0, 1% Triton X-100, and 0.1% SDS), four times with 250 mM RIPA (10 mM Tris-Cl pH 8.0, 250 mM NaCl, 0.1 mM EDTA pH 8.0, 1% Triton X-100, and 0.1% SDS) and twice with TE buffer pH 8.0 (10 mM Tris-Cl pH 8.0 and 0.1 mM EDTA pH 8.0). The samples were subjected to proteinase K treatment in a 50 μl of TE buffer pH 8.0 with 5 μl of 20 mg/ml of proteinase K. The proteinase K was heat inactivated for 95 °C for 5 min in a 100 μl reaction with 1X NEBNext High-Fidelity PCR Mix, during PCR with primers containing molecular indices (listed in Table 1) with the following program; 72°C for 3min, 95°C for 5min, {98°C for 10 sec, 63°C for 30 sec, 72°C for 30 sec} for 12 cycles, 72°C for 5 min, and hold at 4°C. The PCR reaction was purified with bead-based size selection to remove fragments larger than 1000 bp. Ampure Xp beads were added to the PCR reaction in a ratio of 0.2X ratio to bind larger fragments. The magnetic beads were separated with the help of magnetic rack and supernatant was transferred to a fresh tube. Ampure Xp beads were added to the PCR reaction in a ratio of 0.8X to bind the target library. After PCR purification, libraries were analyzed on Agilent Bioanalyzer for size distribution and the concentration was measured using a Qubit fluorometer. The finished libraries were pooled in equimolar amounts and sequenced on Illumina NextSeq 500.

### TAF-ChIP and Conventional ChIP qPCR

2 μl of TAF as well as conventional ChIP library was used for checking enrichment with various primer pairs (listed in Table 1) on Applied Biosystem ViiA™ 7 real time machine using SYBR green reagent (Life technologies, Cat No-4367659).

### Demultiplexing and mapping

De-multiplexing and fastq file conversion were performed using blc2fastq (v.1.8.4). Reads from ChIP-Seq libraries were mapped using bowtie2 (v. 2.2.8) [23], and filtered for uniquely mapped reads. The genome build and annotation used for all *Drosophila* samples was BDGP6 (ENSEMBL release 84). The genome build and annotation used for the K562 samples was hg38 (ENSEMBL release 84).

### Normalization, peak calling and overlaps

The mapped BAM files were normalized to RPKMs using deepTools, and bigwig coverage files were generated. Peak calling was performed using MACS2 (v 2.1.1-20160309) [24]. The peaks were called with following settings: (A) For *Drosophila* H3K4Me3; macs2 callpeak -t ChIP.bam -c Control.bam -f BAMPE -g dm –q 0.05. (B) For *Drosophila* H3K9Me3; macs2 callpeak -t ChIP.bam -c Control.bam -f BAMPE -g dm --broad --broad-cutoff 0.05. (C) For K562 H3K9Me3 and H3K27Me3; macs2 callpeak -t ChIP.bam -c Control.bam -f BAMPE -g hs --broad --broad-cutoff 0.05. The resulting peaks were annotated with the ChIPseeker package from Bioconductor, using nearest gene to peak summit as assignment criteria [25]. The individual peaks of corresponding modification and approach were merged using bedTools, except ENCODE peak files [26]. For ENCODE, owing to high variability between the replicates we used replicated peaks provided by ENCODE database. After merging, the overlaps were calculated with ChIPpeakAnno package with following commands; (A) For H3K4Me3 in *Drosphila* NSCs; overlap = findOverlapsOfPeaks(ConvK4Me3, TAFK4Me3, maxgap = 100), (B) For H3K9Me3 in *Drosphila*; overlap = findOverlapsOfPeaks(ConvK9Me3, TAFK9Me3, maxgap = 200), (C) For H3K27Me3 in K562; overlap = findOverlapsOfPeaks(ENCODEK27Me3, TAFK27Me3, maxgap = 4000), and (D) For H3K9Me3 in K562; overlap = findOverlapsOfPeaks(ENCODEK9Me3, TAFK9Me3, maxgap = 4000).

### Computational Scripts

All the parameters used for computational analysis and detailed scripts are provided in a separate supplementary text file. The heatmaps were generated using deepTools (v 3.5.1) [27].

---

**Supplementary Figure 1.**
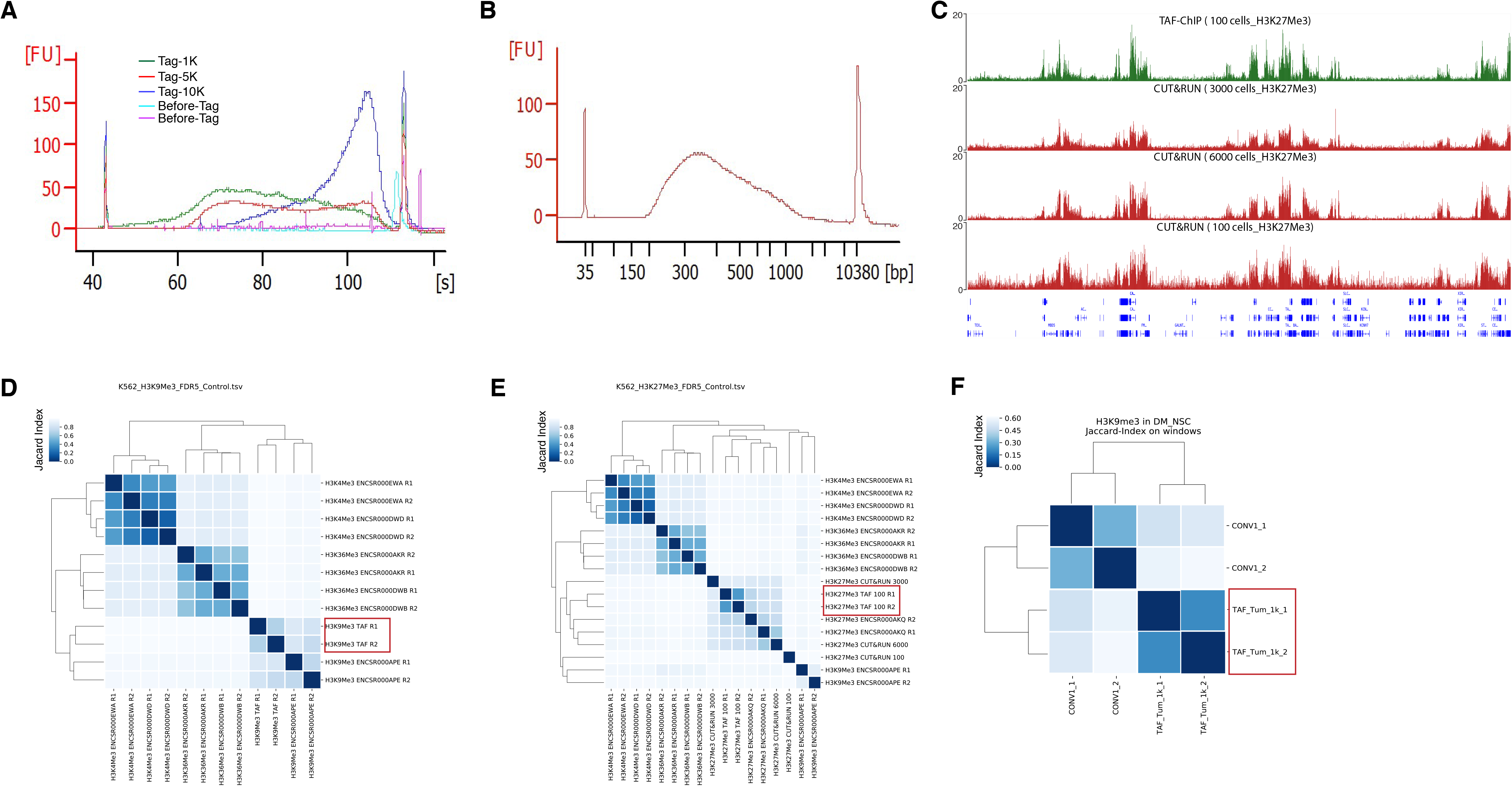
Comparison of TAF-ChIP with ENCODE, conventional ChIP-Seq, and CUT&RUN. **(A)** Bioanalyzer profile before and after tagmentation with indicated cell numbers. The amount of Tn5 tagmentase was kept constant in all conditions. **(B)** Bioanalyzer profile of a representative TAF-ChIP library showing the size distribution of final library fragments. **(C)** Genome browser track example of H3K27Me3 with TAF-ChIP approach and recently published CUT&RUN method with different cell numbers, as indicated in the labels. The label below the tracks shows the gene model, and the y-axis represents normalized read density in reads per million (rpm). **(D)** Clusterogram of TAF-ChIP H3K9Me3 (highlighted with a rectangular box) and indicated datasets derived from the signal in the peak file. H3K4Me3 and H3K36Me3 were included as controls. Note that the TAF-ChIP replicates for H3K9Me3 cluster together with the equivalent datasets from ENCODE. The legend in the left indicates the distance based on Jaccard Index. **(E)** Clusterogram of TAF-ChIP H3K27Me3 datasets (highlighted by a rectangular box) with equivalent datasets from ENCODE, CUT&RUN datasets (from 100, 3000, and 6000 cells), and non-related histone modification datasets used as control. **(F)** Clusterogram of TAF-ChIP H3K9Me3 datasets (highlighted by a rectangular box) with conventional ChIP-Seq dataset from *Drosophila* NSCs.

**Supplementary Figure 2.**
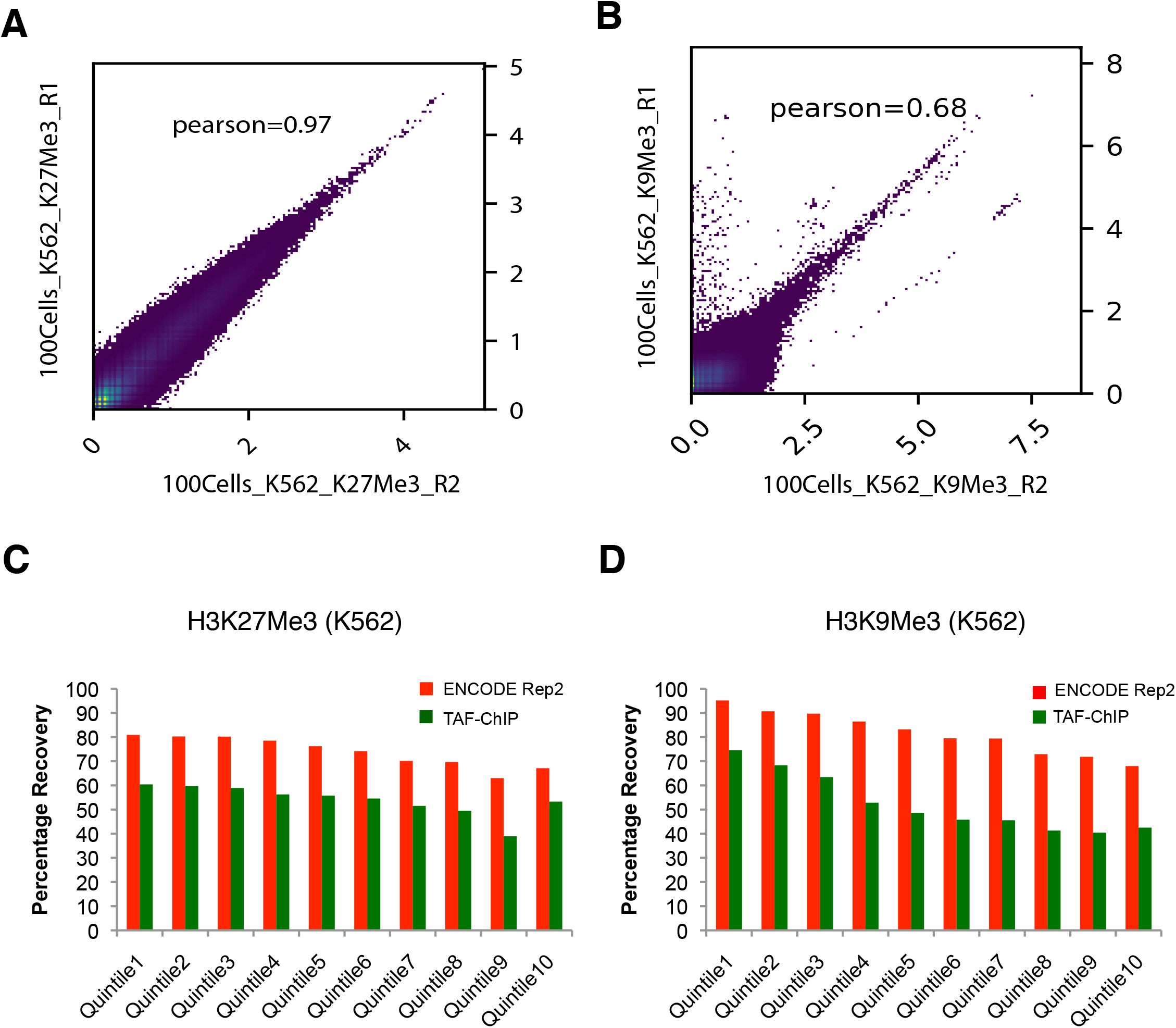
TAF-ChIP libraries evaluated in terms of replicate variability and peaks recovery, from 100 K562 cells. **(A, B)** Pearson correlation between the indicated replicates of TAF-ChIP samples from 100 K562 cells, across equal sized bins of 2000 bp. **(C, D)** Percentage recovery of peaks in TAF-ChIP dataset compared to a corresponding ENCODE ChIP-Seq dataset. The quantiles were sorted according to increasing FDR (provided by MACS2). Recovery of peaks of one of the replicates of the ENCODE ChIP-Seq dataset when the other is used as reference is shown for comparison. For H3K27Me3 **(C)** and H3K9Me3 **(D)** histone modifications in K562 cells.

**Supplementary Figure 3.**
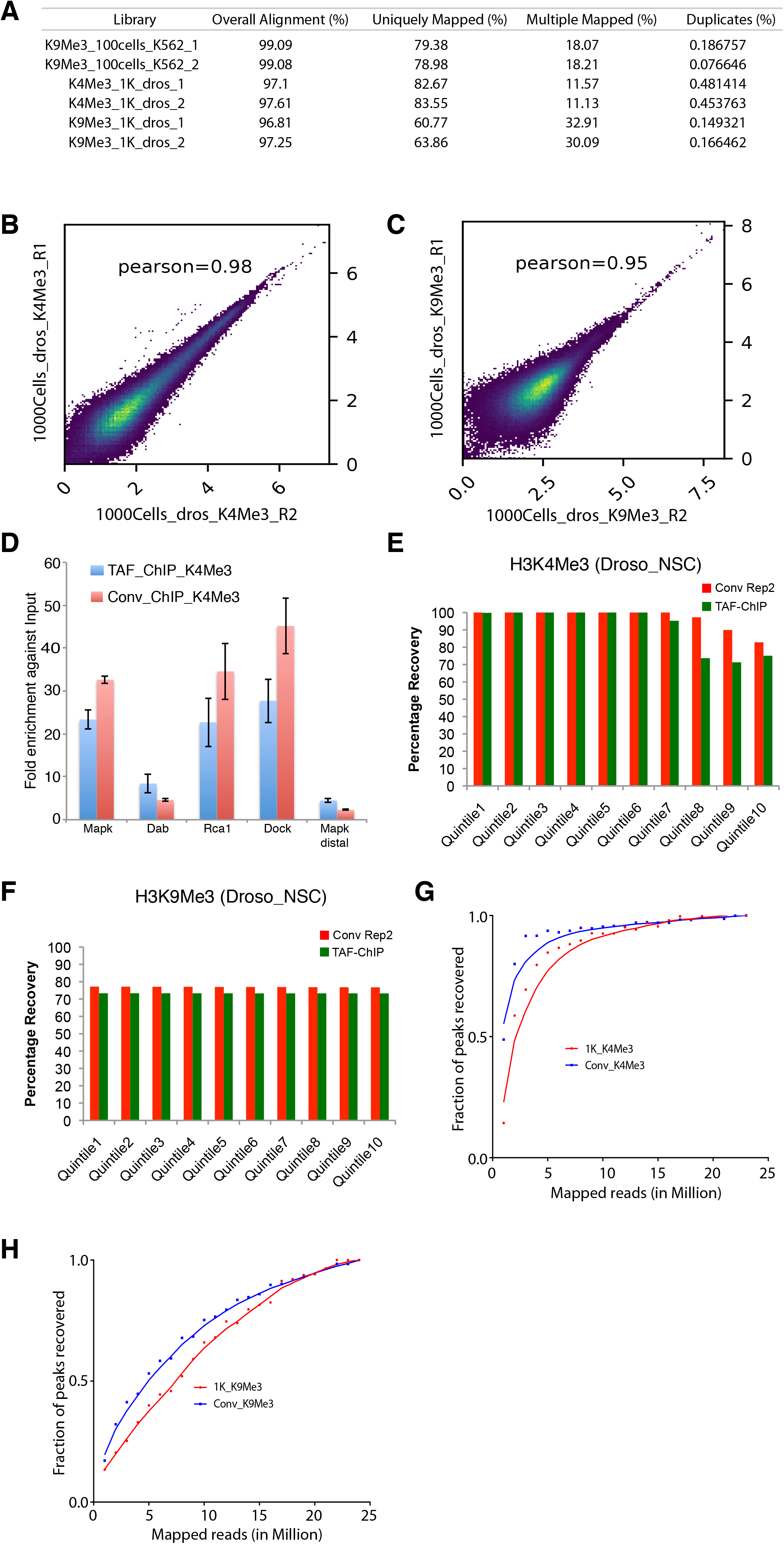
TAF-ChIP libraries evaluated in terms of replicate variability and peaks recovery, from 1000 *Drosophila* NSCs. **(A)** Table showing the percentage of mapped reads from TAF-ChIP experiments. The read duplicates were quantified using Picard Tools. **(B, C)** Pearson correlation between the indicated replicates of TAF-ChIP samples from Drosophila 1000 cells, across equal sized bins of 500 bp. **(D)** ChIP-qPCR analysis of H3K4Me3 enrichment at the tested loci in TAF-ChIP and conventional approach against respective input controls, fragmented DNA in conventional and H3 for TAF. The tested loci Mapk, Dab, Rca1, and Dock showed enrichment in conventional and TAF-ChIP Seq results while Mapk distal did not show any enrichment in either approach. **(E, F)** Percentage recovery of peaks in TAF-ChIP datasets compared to conventional ChIP-Seq datasets. The quantiles were sorted according to increasing FDR (provided by MACS2). Recovery of peaks of one of the replicates of conventional ChIP-Seq using the other as reference is shown for comparison. For H3K4Me3 **(E)** and H3K9Me3 **(F)** histone modifications in *Drosophila* NSCs. **(G, H)** Fraction of peaks recovered for H3K4Me3 (G) and H3K9Me3 (H) TAF-ChIP samples from *Drosophila*, at various sequencing depths.

**Supplementary Figure 4.**
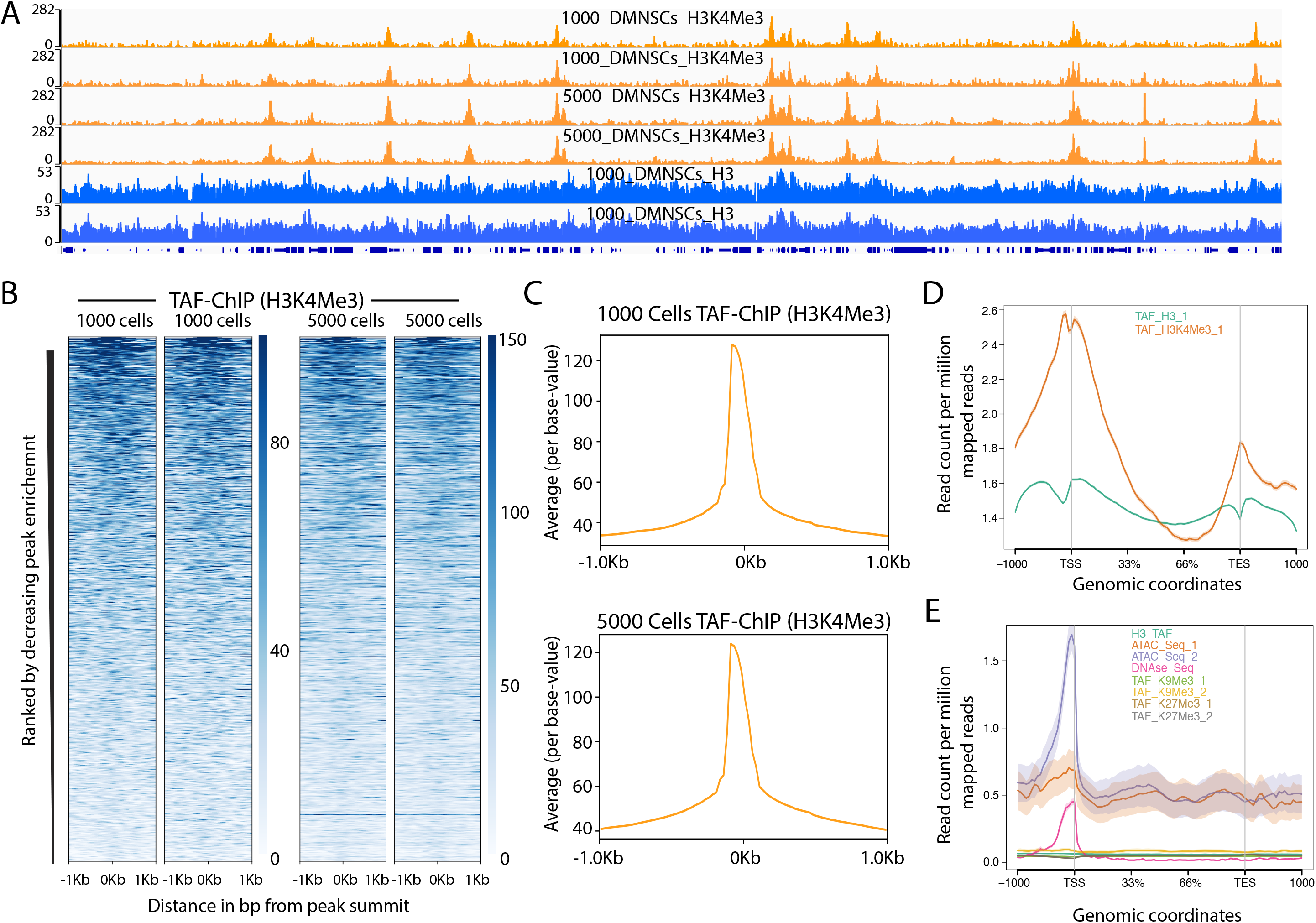
TAF-ChIP results from different amounts of starting material are identical and H3 TAF-ChIP do not show any visible biases for open chromatin. **(A)** Genome browser track example of H3K4Me3 ChIP performed for 2 replicates in 1000 and 5000 FACS sorted *Drosophila* NSCs along with H3 controls, as indicated in the labels. The label below the tracks shows the gene model, and the y-axis represents normalized read density in reads per million (rpm). **(B)** Heatmaps of TAF-ChIP datasets for H3K4Me3 from 1000 and 5000 sorted *Drosophila* NSCs. Rows indicate all the peaks and are sorted by decreasing affinities in the conventional ChIP-Seq data sets. The color labels to the right indicate the level of enrichment. **(C)** Average profile of TAF-ChIP data from 1000 and 5000 NSCs, centered at the peaks for the H3K4Me3 modification. The y-axis depicts average per base peak signal while the x-axis depicts genomic coordinates centered at the peaks. **(D)** Metagene profiles of H3 and H3K4Me3 from 1000 sorted *Drosophila* NSCs with standard error to the mean for all genes, −1000bp upstream of transcription start sites (TSSs) and +1000bp downstream of transcription end sites (TESs). Read counts per million of mapped reads is shown on the y-axis, while the x-axis depicts genomic coordinates. **(E)** Metagene profiles of indicated datasets with standard error to the mean for all genes in K562 cells, - 1000bp upstream of transcription start sites (TSSs) and +1000bp downstream of transcription end sites (TESs). Read counts per million of mapped reads is shown on the y-axis, while the x-axis depicts genomic coordinates. The ATAC-Seq and DNAse-Seq datasets are derived from PMID: 26280331.

**Supplementary Figure 5.**
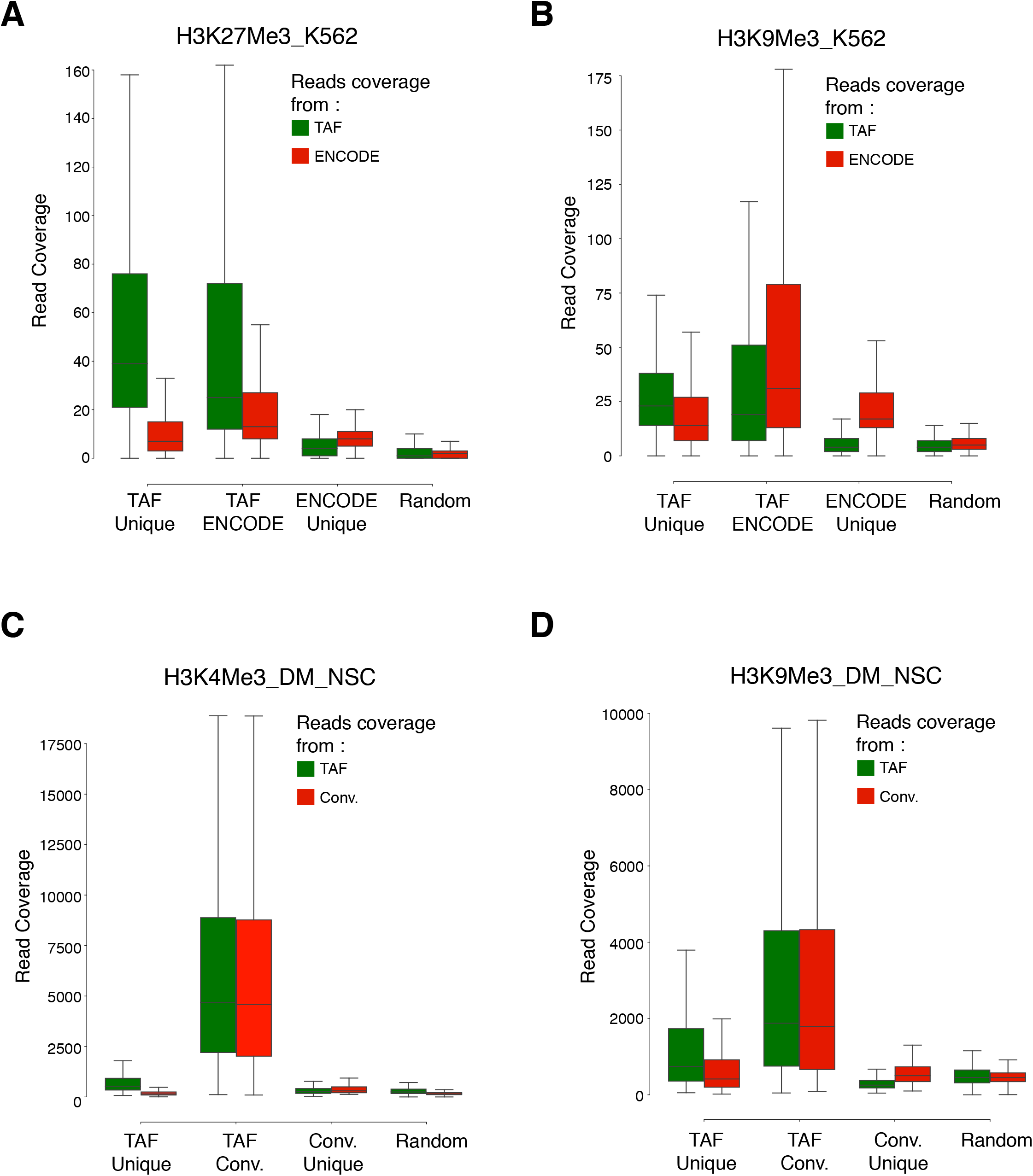
Coverage of reads over the overlapping and unique peaks identified in the TAF-ChIP and conventional datasets. **(A-D)** Boxplots showing coverage of raw reads over the peaks identified either uniquely in TAF (TAF Unique) or conventional ChIP-Seq approach (ENCODE Unique or Conv. Unique), or common to both of the approaches (TAF ENCODE or TAF Conv.). Shown also is the coverage of reads over randomly selected regions of comparable size (Random). From K562 cells for H3K27Me3 **(A)** and H3K9Me3 **(B)**, and from *Drosophila* neural stem cells for H3K4Me3 **(C)** and H3K9Me3 **(D)**.

## References

1. Orlando, V., Mapping chromosomal proteins in vivo by formaldehyde-crosslinked-chromatin immunoprecipitation. Trends Biochem Sci, 2000. 25(3): p. 99–104.

2. Mundade, R., et al., Role of ChIP-seq in the discovery of transcription factor binding sites, differential gene regulation mechanism, epigenetic marks and beyond. Cell Cycle, 2014. 13(18): p. 2847–52.

3. Stathopulos, P.B., et al., Sonication of proteins causes formation of aggregates that resemble amyloid. Protein Sci, 2004. 13(11): p. 3017–27.

4. Mieczkowski, J., et al., MNase titration reveals differences between nucleosome occupancy and chromatin accessibility. Nat Commun, 2016. 7: p. 11485.

5. Gutierrez, G., et al., Subtracting the sequence bias from partially digested MNase-seq data reveals a general contribution of TFIIS to nucleosome positioning. Epigenetics Chromatin, 2017. 10(1): p. 58.

6. Dingwall, C., G.P. Lomonossoff, and R.A. Laskey, High sequence specificity of micrococcal nuclease. Nucleic Acids Res, 1981. 9(12): p. 2659–73.

7. Zheng, X., et al., Low-Cell-Number Epigenome Profiling Aids the Study of Lens Aging and Hematopoiesis. Cell Rep, 2015. 13(7): p. 1505–1518.

8. Adli, M., J. Zhu, and B.E. Bernstein, Genome-wide chromatin maps derived from limited numbers of hematopoietic progenitors. Nat Methods, 2010. 7(8): p. 615–8.

9. van Galen, P., et al., A Multiplexed System for Quantitative Comparisons of Chromatin Landscapes. Mol Cell, 2016. 61(1): p. 170–80.

10. Ross-Innes, C.S., et al., Differential oestrogen receptor binding is associated with clinical outcome in breast cancer. Nature, 2012. 481(7381): p. 389–93.

11. Schmidl, C., et al., ChIPmentation: fast, robust, low-input ChIP-seq for histones and transcription factors. Nat Methods, 2015. 12(10): p. 963–965.

12. Skene, P.J., J.G. Henikoff, and S. Henikoff, Targeted in situ genome-wide profiling with high efficiency for low cell numbers. Nat Protoc, 2018. 13(5): p. 1006–1019.

13. Picelli, S., et al., Tn5 transposase and tagmentation procedures for massively scaled sequencing projects. Genome Res, 2014. 24(12): p. 2033–40.

14. Seguin-Orlando, A., et al., Ligation bias in illumina next-generation DNA libraries: implications for sequencing ancient genomes. PLoS One, 2013. 8(10): p. e78575.

15. Ghavi-Helm, Y., B. Zhao, and E.E. Furlong, Chromatin Immunoprecipitation for Analyzing Transcription Factor Binding and Histone Modifications in Drosophila. Methods Mol Biol, 2016. 1478: p. 263–277.

16. Berger, C., et al., FACS purification and transcriptome analysis of drosophila neural stem cells reveals a role for Klumpfuss in self-renewal. Cell Rep, 2012. 2(2): p. 407–18.

17. Bowman, S.K., et al., The tumor suppressors Brat and Numb regulate transit-amplifying neuroblast lineages in Drosophila. Dev Cell, 2008. 14(4): p. 535–46.

18. Wang, H., et al., Aurora-A acts as a tumor suppressor and regulates self-renewal of Drosophila neuroblasts. Genes Dev, 2006. 20(24): p. 3453–63.

19. Consortium, E.P., An integrated encyclopedia of DNA elements in the human genome. Nature, 2012. 489(7414): p. 57–74.

20. Landt, S.G., et al., ChIP-seq guidelines and practices of the ENCODE and modENCODE consortia. Genome Res, 2012. 22(9): p. 1813–31.

21. Buenrostro, J.D., et al., Transposition of native chromatin for fast and sensitive epigenomic profiling of open chromatin, DNA-binding proteins and nucleosome position. Nat Methods, 2013. 10(12): p. 1213–8.

22. Lihua J Zhu., et al., ChIPpeakAnno: a Bioconductor package to annotate ChIP-seq and ChIP-chip data. BMC Bioinformatics. 2010; 11: 237.

23. Langmead, B., Aligning short sequencing reads with Bowtie. Curr Protoc Bioinformatics, 2010. Chapter 11: p. Unit 11–7.

24. Zhang, Y., et al., Model-based analysis of ChIP-Seq (MACS). Genome Biol, 2008. 9(9): p. R137.

25. Yu, G., L.G. Wang, and Q.Y. He, ChIPseeker: an R/Bioconductor package for ChIP peak annotation, comparison and visualization. Bioinformatics, 2015. 31(14): p. 2382–3.

26. Quinlan AR. BEDTools: the Swiss-army tool for genome feature analysis. Curr Protoc Bioinformatics, 2014; 47: 11.12.1–11.12.34.

27. Ramirez, F., et al., deepTools2: a next generation web server for deep-sequencing data analysis. Nucleic Acids Res, 2016. 44(W1): p. W160–5.

